# Host Genetic Background and Gut Microbiota Contribute to Differential Metabolic Responses to Fructose Consumption in Mice

**DOI:** 10.1101/439786

**Authors:** In Sook Ahn, Jennifer M. Lang, Christine A. Olson, Graciel Diamante, Guanglin Zhang, Zhe Ying, Hyae Ran Byun, Ingrid Cely, Jessica Ding, Peter Cohn, Ira Kurtz, Fernando Gomez-Pinilla, Aldons J. Lusis, Elaine Y. Hsiao, Xia Yang

**Affiliations:** Department of Integrative Biology and Physiology, University of California, Los Angeles, CA; Department of Medicine/Division of Cardiology, David Geffen School of Medicine, University of California, Los Angeles, CA; Department of Medicine, Division of Nephrology, University of California, Los Angeles, CA; Brain Research Institute, University of California, Los Angeles, CA; Department of Neurosurgery, University of California, Los Angeles, CA; Molecular Biology Institute, University of California, Los Angeles, CA; Institute for Quantitative and Computational Biosciences, University of California, Los Angeles, CA

**Author notes:** These authors contributed equally (In Sook Ahn, Jennifer M. Lang). **Corresponding author:** Xia Yang, Ph.D., Department of Integrative Biology and Physiology, University of California, Los Angeles, Los Angeles, CA 90095, USA. Supplemental Figures 1-5 are available from the “Supplementary data” link in the online posting of the article and from the same link in the online table of contents at https://academic.oup.com/jn/. Supplemental Tables 1 and 2 were uploaded separately as excel files at https://figshare.com/s/ff6ffce17c10f6a1eb1f. Abbreviations used: AM, *Akkermansia muciniphila*; AUC, area under the curve; B6, C57BL/6J; DBA, DBA/2J; FDR, false discovery rate; FMT, fecal microbiota transplant; FVB, FVB/NJ; IPGTT, intraperitoneal glucose tolerance test; PCoA, principal coordinates analysis; PERMANOVA, permutational multivariate analysis of variance; TH, tyrosine hydrogenase.

**Keywords:** gut microbiota, fructose, metabolic syndrome, fecal transplant, *Akkermansia*, microbiota-host interaction, gene by diet interaction

## Abstract

**Background:** It is unclear how high fructose consumption induces disparate metabolic responses in genetically diverse mouse strains.

**Objective:** We aim to investigate whether the gut microbiota contributes to differential metabolic responses to fructose.

**Methods:** Eight-week-old male C57BL/6J (B6), DBA/2J (DBA), and FVB/NJ (FVB) mice were given 8% fructose solution or regular water (control) for 12 weeks. The gut microbiota composition in cecum and feces was analyzed using 16S rDNA sequencing, and PERMANOVA was used to compare community across mouse strains, treatments, and time points. Microbiota abundance was correlated with metabolic phenotypes and host gene expression in hypothalamus, liver and adipose tissues using Biweight midcorrelation. To test the causal role of the gut microbiota in determining fructose response, we conducted fecal transplants from B6 to DBA mice and vice versa for 4 weeks, as well as gavaged antibiotic-treated DBA mice with *Akkermansia* for 9 weeks, accompanied with or without fructose treatment.

**Results:** Compared to B6 and FVB, DBA mice had significantly higher Firmicutes/Bacteroidetes ratio and lower baseline levels of *Akkermansia* and S24-7 (*P* < 0.05), accompanied by metabolic dysregulation after fructose consumption. Fructose altered specific microbial taxa in individual mouse strains, such as a 7.27-fold increase in *Akkermansia* in B6 and 0.374-fold change in Rikenellaceae in DBA (FDR < 5%), which demonstrated strain-specific correlations with host metabolic and transcriptomic phenotypes. Fecal transplant experiments indicated that B6 microbes conferred resistance to fructose-induced weight gain in DBA mice (*F* = 43.1, *P* < 0.001), and *Akkermansia* colonization abrogated the fructose-induced weight gain (*F* = 17.8, *P* <0.001) and glycemic dysfunctions (*F* = 11.8, *P* = 0.004) in DBA mice.

**Conclusions:** Our findings support that differential microbiota composition between mouse strains is partially responsible for host metabolic sensitivity to fructose, and that *Akkermansia* is a key bacterium that confers resistance to fructose-induced metabolic dysregulation.

## Introduction

The drastic increase in fructose consumption over the past few decades has been paralleled by the rising prevalence of metabolic syndrome and diabetes (1). Our recent systems nutrigenomics study unraveled the impact of fructose on transcriptome, epigenome and gene-gene interactions in the hypothalamus, which is the main regulator of metabolism (2). Additionally, studies on the liver and adipose tissues have also revealed fructose-induced alterations in genes involved in several aspects of systemic metabolism (3–5). Interestingly, genetically diverse mouse strains demonstrate disparate metabolic responses to fructose consumption, a finding reproducible across multiple studies, including ours (5,6). However, the causal mechanisms underlying the inter-individual differences in metabolic responses to fructose have yet to be elucidated.

The gut microbiome is emerging as an important modulator of metabolism, obesity, and other metabolic disorders (7). Its dynamic nature, ease of manipulation, and response to dietary intervention have made the gut microbiome a suitable therapeutic target to mitigate metabolic syndrome (8). Diet influences trillions of gut microorganisms as early as one day post dietary intervention (9). The gut microbiome has been suggested to contribute to phenotypic diversity in different mouse strains on a high-fat high-sucrose diet (10,11). It is plausible that the gut microbiome also plays a role in determining inter-individual variability in metabolic responses to fructose consumption.

Dietary fructose is mainly absorbed in the small intestine, where it can be extensively metabolized into glucose and organic acids. However, excessive amount of fructose that is not absorbed in the small intestine due to malabsorption or high intake can undergo colonic bacterial fermentation, resulting in the production of metabolites such as short-chain fatty acids (12). Fructose also has the potential to modify the microbiota community and its normal function. For instance, fructose metabolites can shape the gut environment and be an energy source for the gut microbiota (13). Fructose can also suppress gut bacterial colonization by silencing a colonization factor in a commensal bacterium (14).

Here, we investigated gut microbiota as a potential link between fructose consumption and differential metabolic phenotypes in mice with diverse genetic backgrounds. We tested our paradigm in three mouse strains, namely C57BL/6J (B6), DBA/2J (DBA), and FVB/NJ (FVB), which have contrasting metabolic susceptibility to high caloric diets (5,6,11).

## Methods

### Animals and study design

To investigate the role of the gut microbiota in host responses to fructose consumption, we examined microbial composition in the context of differential metabolic and transcriptomic responses in multiple mouse strains (**Supplemental Figure 1**). Seven-week-old male mice (20-25g) from three inbred strains, namely B6, DBA, and FVB, were obtained from the Jackson Laboratory (Bar Harbor, ME) and housed in a pathogen-free barrier facility with a 12-hour light/dark cycle at University of California, Los Angeles. Mice were fed Lab Rodent Diet 5001, containing 234 g/kg protein, 45 g/kg fat, 499 g/kg carbohydrate, and 2.89 kcal/g metabolizable energy (LabDiet, St. Louis, MO). After a one-week acclimation period, mice from each strain were randomly divided into two groups. One group was provided with regular water (control group, *n*=8-10 mice/strain) and the other group was given 8% (w/v) fructose (3.75 kcal/g energy; NOW Real Food, Bloomingdale, IL) dissolved in regular water (fructose group, *n*=10-12 mice/strain) for 12 weeks *ad libitum*. Food and drink intake were monitored daily on a per-cage basis. Distinct alterations in body weight, fat mass, plasma lipids, glycemic traits, intraperitoneal glucose tolerance test (IPGTT), and the transcriptome of liver, adipose, and hypothalamus in response to fructose consumption across mouse strains were observed and previously published (5). Gut microbiota composition of cecal and fecal samples were analyzed. Fructose-responsive microbiota were correlated with metabolic phenotypes and host gene expression to prioritize microbial taxa that may contribute to differential host fructose responses. Lastly, fecal microbiota transplant (FMT) and *Akkermansia muciniphila* (AM) colonization experiments were conducted to test the causal role of gut microbiota in determining mouse strain-specific fructose responses. The study was performed in accordance with National Institutes of Health Guide for the Care and Use of Laboratory Animals. All experimental protocols were approved by the Institutional Animal Care and Use Committee at the University of California, Los Angeles.

### Fecal and cecal microbiota analysis

Feces were collected at 1, 2, 4, and 12 weeks of fructose treatment, and cecal contents were collected at the end of fructose treatment (12 weeks). Samples were snap frozen and then stored at −80°C until DNA isolation. Microbial DNA was isolated from samples using the MO BIO PowerSoil®-htp 96 Well Soil DNA Isolation Kit (MO BIO, Carlsbad, CA). For the fecal and cecal samples, the same lot isolation kit was used; for the fecal transplant samples a different lot kit was used. The V4 region (15,16) of the 16S rDNA gene was amplified in triplicate with barcoded primers (515f and 806r) (17). PCR products were quantified with Quant-iTTM PicoGreen® dsDNA Assay Kit (Thermo Fisher Scientific) and all samples from a specific experiment (e.g., fecal, cecal, or fecal transplant experiment) were combined in equal amounts (~250 ng per sample) into a single pooled submission to be purified with the UltraClean PCR® Clean-Up Kit (MO BIO). Single-end reads were generated on the Illumina HiSeq 2500 platform using two lanes for each pool of mixed samples to create an unbiased sequencing approach. Roughly 500 samples were sequenced per submission, and each dataset of fecal, cecal, and fecal transplant samples were sequenced in different submissions. Raw sequences were processed using QIIME to produce de-multiplexed and quality-controlled sequences. Reads were binned into OTUs at 97% similarity using UCLUST against the Greengenes reference database (18). Singletons, OTUs representing less than 0.005% total relative abundance, unsuccessfully sequenced samples, and outliers were removed. Samples were normalized to a rarefied level specific for each dataset to reduce the effect of unequal sequencing depth to result in 60,000, 106,815, and 37,614 reads per sample for fecal, cecal, and fecal transplant samples, respectively. A total of 180 fecal, 46 cecal, and 170 fecal transplant samples were used for downstream analysis.

### Correlation analysis between gut microbiota and metabolic phenotypes or fructose signature genes

Correlation analysis was performed between the relative abundance or proportion of microbiota with metabolic phenotypes including body weight, adiposity, and area under the curve (AUC) for glucose tolerance. Microbiota proportions were also correlated with differentially expressed genes in response to fructose consumption, or “fructose signature genes”, in hypothalamus, liver, and adipose tissue. The fructose signature genes correlated with bacterial abundance were classified for their biological functions in Gene Ontology, REACTOME, and KEGG using the GSEA tool (19).

### Antibiotics treatment prior to fecal transplant or *Akkermansia* colonization

Six-week-old B6 or DBA recipient mice were orally gavaged with a solution of vancomycin (50 mg/kg), neomycin (100 mg/kg), and metronidazole (100 mg/kg) twice daily for 7 days. Ampicillin (1 mg/mL) was also provided *ad libitum* in drinking water (20). Antibiotic-treated mice were housed in sterile cages with sterile water and food throughout the experiment. Following antibiotics treatment, mice were used in either fecal transplant or *Akkermansia* colonization experiments, as described below.

### Fecal transplant between B6 and DBA mice

Reciprocal fecal transplant between B6 and DBA mice was conducted. B6 mice receiving DBA feces were designated as B6(DBA) and DBA mice receiving B6 feces were designated as DBA(B6). To serve as control groups, DBA mice were transplanted with DBA feces and designated as DBA(DBA), and B6 mice were transplanted with B6 feces and designated as B6(B6). Fecal transplant was performed according to previous studies with some modification (21,22). Briefly, freshly collected feces were pooled from 4 donor mice and suspended at 40mg/mL concentration in anaerobic PBS. The suspension (150 *μ*L) was orally gavaged to the recipient mice for 4 weeks. Total time between collecting feces and delivery of microbial contents into recipients was kept as short as possible (< 15 min) to protect anaerobes. After 1 week of fecal transplant, mice from each of the four transplant experiments, namely B6(DBA), B6(B6), DBA(B6), DBA(DBA), were divided into two groups and treated with 8% fructose or regular water for 12 weeks (*n* = 7-14 mice/group). Body weight and IPGTT were measured using the methods described previously (5).

### *Akkermansia* colonization in DBA mice

*Akkermansia* colonization was conducted according to Olson et al. (20). Antibiotic-treated DBA mice were orally gavaged with 200 *μ*L bacterial suspension (5 × 10^9^ cfu/mL in anaerobic PBS) throughout the experiment. After 1 week of bacterial gavage, DBA mice were treated with 8% fructose or regular water for 8 weeks (*n* = 10-14/group). DBA mice receiving anaerobic PBS served as control (*n* = 8-10/group). Body weight and IPGTT were measured as described previously (5).

### Statistical analysis

The microbiota data was summarized into relative abundance by taxonomic level in QIIME, and communities were visualized with principal coordinates analysis (PCoA) based on the weighted UniFrac distance measure (23). Categorical groups (treatment, time, mouse strain) were confirmed to have similar multivariate homogeneity of group dispersions to allow them to be compared using the non-parametric PERMANOVA (permutational multivariate analysis of variance) test with the adonis function (24). Microbial composition was analyzed at the phylum, family, and genus taxonomic levels using the Statistical Analysis of Metagenomic Profiles (STAMP) software (25).

Linear discriminant analysis (LDA) effect size (LEfSe) was used to identify taxa differentially represented between three mouse strains using standard parameters (*P* < 0.05, LDA score > 2.0) (26). Using the identified features from the LEfSe analysis, we selected six fecal genera which showed contrasting patterns in their baseline levels between DBA and the other two mouse strains, B6 and FVB. To visualize the baseline differences of these six genera between the three mouse strains, boxplots were plotted using centered log-ratio (CLR) transformed values from OTU counts with the ‘rgr’ package in R software (24). The difference between strains was assessed using a one-way ANOVA followed by Sidak’s post hoc test. Because *Turicibacter* was not normally distributed, the difference between strains was analyzed by Kruskal-Wallis test followed by Dunn’s test.

Taxa that differed between the fructose and regular water groups were identified using White’s non-parametric T-test (27), followed by Storey’s false discovery rate (FDR) estimation using relative abundance data and the Statistical Analysis of Metagenomic Profiles (28). A one-way ANOVA followed by Sidak’s post hoc test was used to determine the difference in Firmicutes/Bacteroidetes (F/B) ratio between three mouse strains. F/B ratio was calculated by dividing the proportion of Firmicutes and Bacteroidetes for each sample, and then log-transformed to achieve normal distribution.

Correlation between gut microbiota and metabolic phenotypes or fructose signature genes from individual tissues, was assessed using Biweight midcorrelation (bicor) (29). Statistical *P*-values were adjusted using the Benjamini-Hochberg approach and FDR < 0.05 was considered significant.

To analyze body weight gain and glucose tolerance data involving multiple time points in fecal transplant and *Akkermansia* colonization experiments, a three-way repeated-measures ANOVA followed by Sidak’s post hoc test was used. The effects and interaction of three factors (microbial manipulation, fructose, time) on the metabolic phenotypes were tested.

To evaluate the effects of fructose under each unique microbiota manipulation (e.g., within *Akkermansia*-colonized groups or within B6 mice transplanted with DBA microbe) across multiple time points, a two-way repeated-measures ANOVA was used. Fructose treatment was used as the between-group factor and time was used as the within-subject factor. To assess the effect of a given microbiota manipulation under fructose treatment across multiple time points, a two-way repeated-measures ANOVA was used. Microbiota manipulation was used as the between-group factor and time was used as the within-subject factor. If the interaction was significant, microbiota manipulations within each time point were compared by Sidak’s post hoc test. Statistical analyses were performed using STATISTICA (version 7, Statsoft Inc., Tulsa, OK) and GraphPad Prism (version 8, GraphPad Software Inc., San Diego, CA). Data were expressed as means ± SEM. *P* < 0.05 was considered statistically significant.

## Results

### Strain-specific metabolic responses to fructose treatment

We have previously reported that B6, DBA, and FVB mice demonstrated striking differences in their metabolic responses to 8% fructose treatment for 12 weeks (5). DBA mice were more sensitive to fructose in terms of obesity and diabetes-related phenotypes including body weight, adiposity, and glucose intolerance. B6 mice showed significant increases while FVB mice had decreases in plasma cholesterol levels. These results demonstrate strong inter-strain variability in fructose response in genetically divergent mouse strains. Importantly, the disparate metabolic responses were not due to differences in overall energy intake [14.6 ± 0.384, 17.9 ± 0.880, or 18.0 ± 0.403 kcal/(mouse · d) for B6, DBA, or FVB, respectively] (5). The differences in metabolic responses (DBA > B6 or FVB) were also not correlated with the amount of fructose water intake [FVB (23.6 ± 1.36 mL/(mouse · d) > B6 (8.74 ± 0.187 mL/(mouse · d) or DBA (8.47 ± 0.390 mL/(mouse · d)] (5). Amount of intake was likely driven by differences in fructose perception and preference between the mouse strains (6,30).

### Overall effects of fructose on gut microbiota community

The gut microbiota in the three mouse strains was assessed using 16S rDNA sequencing. PCoA plots showed distinct clusters of mouse strain for both the cecal (*P* < 0.001 by PERMANOVA, **Figure 1A**) and the fecal samples (*P* < 0.001, Figure 1B). Time was also a significant factor in fecal samples (*P* < 0.001), with week 12 separating from the earlier weeks when evaluating all mouse strains together (Figure 1C) or each mouse strain separately (Figure 1D-F).

**Figure 1.**
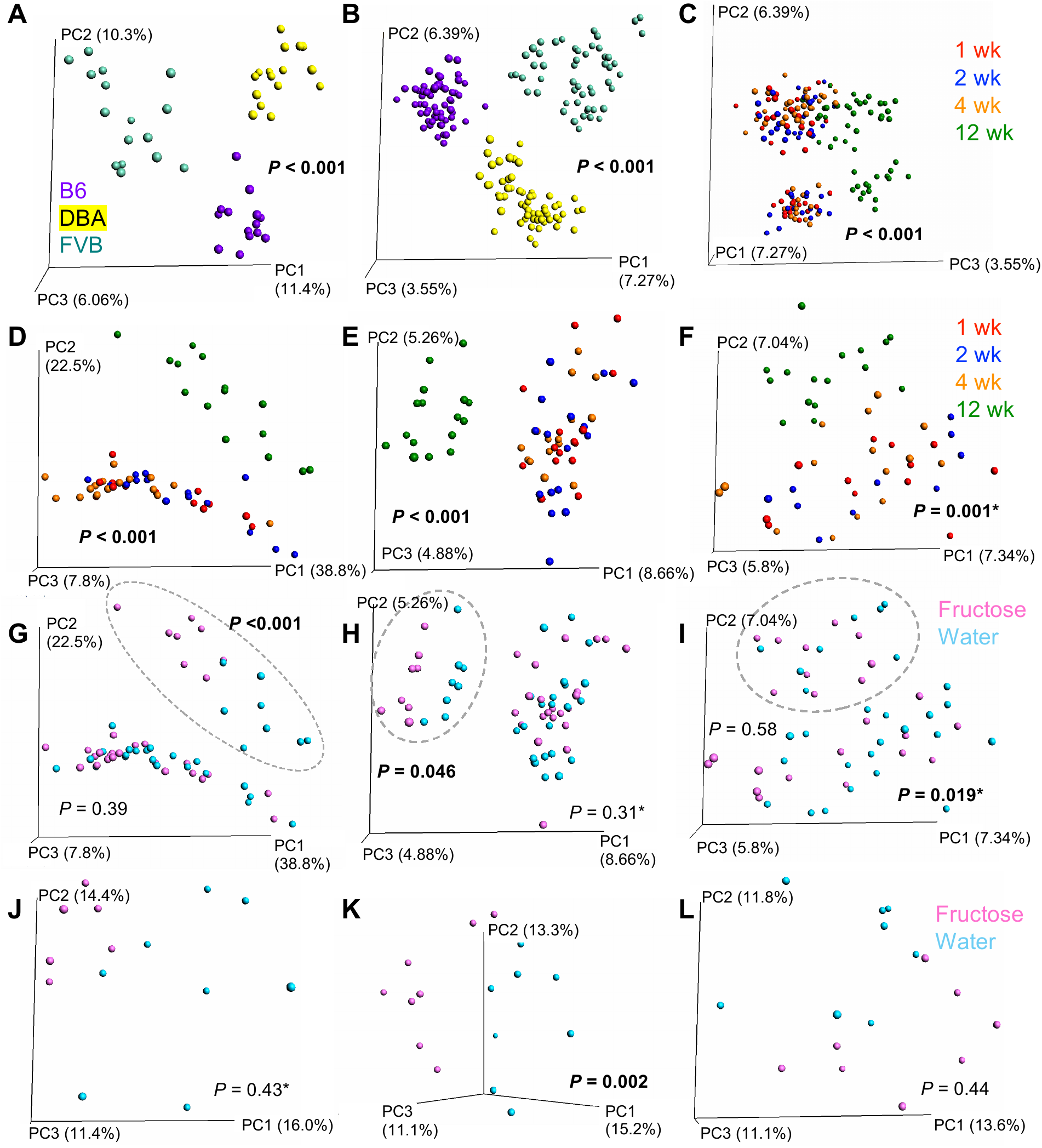
Principal coordinates analysis (PCoA) of cecal and fecal microbiota of B6, DBA, and FVB mice consuming fructose or regular water. (A) Cecal microbiota samples across three mouse strains were shown to separate by strain (A; *n* = 16/strain; week 12). (B-C) Fecal microbiota samples across three mouse strains were separated by both strain (B; *n* = 64/strain across 4 time points) and time (C; n = 16/time point for each strain). Plot C used the same ordination as plot B, except principal coordinate 3 (PC3) was presented as the x-axis to show the relationship with time. (D-F) For each mouse strain, fecal samples were colored by time points for B6 (D), DBA (E), and FVB (F) to show the time effect. (G-I) Fecal samples were colored with fructose or water treatment for B6 (G), DBA (H), and FVB (I) to show treatment effect. Samples for the 12-week time point were shown in dotted circles, with the corresponding *P*-values for fructose treatment effect reported. (J-L) Cecal samples were colored by the fructose or water treatment for B6 (J), DBA (K), and FVB (L). *P-*values were generated by PERMANOVA, and significant results were presented in bold. *P*-values with an asterisk designate that significantly different dispersions were observed, which may influence the *P*-values reported as PERMANOVA assumes similar dispersion.

Fecal microbiota differences were seen between fructose and water groups at 12-weeks in B6 (*P* < 0.001 by PERMANOVA, Figure 1G) and DBA mice (*P* = 0.046, Figure 1H), but not in FVB (*P* = 0.58, Figure 1I). In cecum, the fructose group showed separation from controls for B6 mice (Figure 1J), however, the separation was not statistically significant with PERMANOVA (*P* = 0.43). This result may be influenced by the difference in multivariate spread (*P* = 0.038) since PERMANOVA assumes equal dispersion. Fructose treatment was a significant factor for cecal microbiota composition in DBA (*P* = 0.002, Figure 1K), but not for FVB (*P* = 0.44, Figure 1L). Overall, consistent with the differential effects of fructose on metabolic phenotypes, DBA gut microbiota in both feces and cecum were sensitive to fructose treatment.

### Baseline differences in gut microbiota between mouse strains show correlation with host metabolic phenotypes

Differences in baseline microbial composition can drive distinct host responses to the same dietary manipulation (31,32). At the phylum level, there were no significant differences in cecum Firmicutes/Bacteroidetes (F/B) ratios between the three mouse strains (**Figure 2A**). However, in fecal samples, DBA had significantly higher F/B ratio (1.78 ± 0.169) than B6 (0.346 ± 0.024, *P* < 0.001 by one-way ANOVA) or FVB (0.898 ± 0.091, *P* < 0.001) (Figure 2B), agreeing with the known association of higher Firmicutes abundance with obesity (33) and the increased adiposity in DBA (5). The microbial taxa that accounted for the greatest differences between the three mouse strains included 21 cecal (**Supplemental Figure 2**A, B) and 14 fecal (Supplemental Figure 2C, D) microbial genera based on the LEfSe analysis.

**Figure 2.**
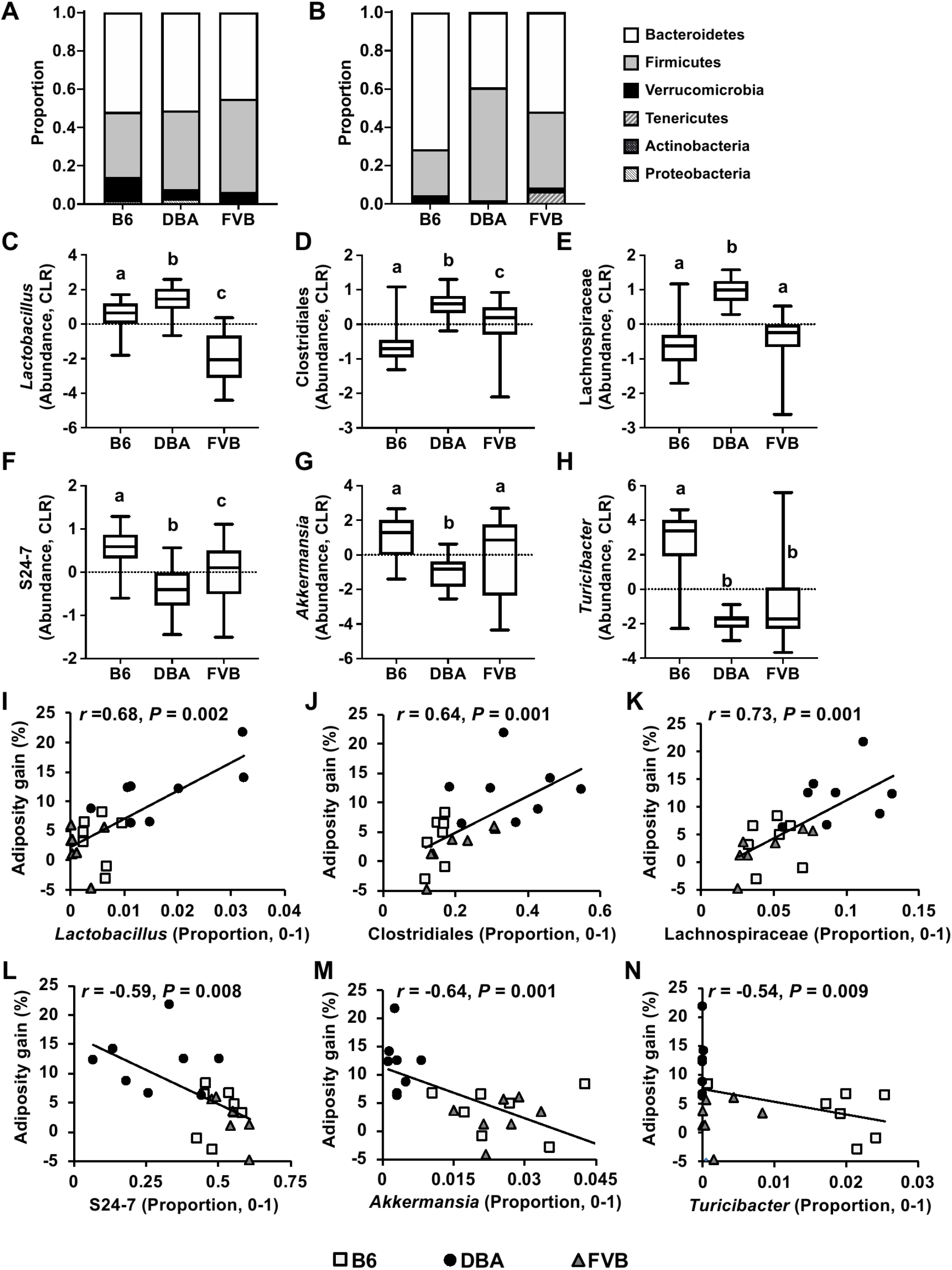
Cecal and fecal baseline microbial composition in B6, DBA, and FVB mice and correlation with adiposity gain. (A-B) Taxa bar plots of baseline cecal (A) and fecal (B) microbiota of three mouse strains at the phylum level. (C-H) Baseline abundance profiles for specific fecal microbiota of three mouse strains at the genus level. Centered log-ratio (CLR) values were used for plotting the abundance of each microbiota. Box and whiskers plots from minimum to maximum showing abundance distribution of *Lactobacillus* (C), unknown genus of Clostridiales (D), unknown genus of Lachnospiraceae (E), unknown genus of S24-7 (F), *Akkermansia (*G), and *Turicibacter* (H). The center line in the box denotes the median value. One-way ANOVA followed by Sidak’s post hoc test was conducted to calculate significant differences between three mouse strains. Labeled means without a common letter differ, *P* < 0.05. *n* = 7-8/group. (I-N) Correlation analysis plots between microbiota baseline proportion and adiposity gain at week 12 of fructose treatment. *r* = Biweight midcorrelation (*bicor*) coefficient, *P* = Benjamini-Hochberg adjusted *P*-values. *n* = 7-8/mouse strain.

We reasoned that if any specific microbial taxon determines the differential fructose response between the mouse strains, its abundance likely shows contrasting patterns between the susceptible strain DBA and the two resistant strains. Out of 21 cecal genera (Supplemental Figure 2B), none showed contrasting patterns between DBA and the two resistant strains. Out of the 14 fecal genera (Supplemental Figure 2D), 6 showed distinct patterns in DBA mice compared to the resistant strains (Figure 2C-H). DBA mice showed higher centered log-ratio (CLR) abundance for *Lactobacillus*, an unknown bacteria of order Clostridiales, and an unknown bacteria of family Lachnospiraceae compared to B6 and FVB. On the other hand, DBA mice had lower CLR abundance for an unknown bacteria of family S24-7, *Akkerman*sia, and *Turicibacter*. To understand the potential role of these microbial taxa in metabolic regulation, the correlations between the proportion of these taxa and adiposity gain across all mouse strains were tested. We found that all taxa were significantly correlated with adiposity gain (*P* < 0.01, Figure 2I-N).

### Fructose-responsive microbiota and correlation with host metabolic phenotypes

Next, we explored the differentially abundant microbiota between fructose and water groups in the three mouse strains at 12 weeks. We observed more fructose responsive microbiota in feces (9 taxa) compared to cecum samples (1 taxon) across mouse strains at the family level (**Table 1**).

In feces, fructose treatment showed the highest impact in B6 mice by altering the abundance of many microbiota (nine families and three genera). Significant decreases were observed in five taxa belonging to phylum Firmicutes. There were also significant increases in S24-7 of phylum Bacteroidetes and Verrucomicrobiaceae. Within the Verrucomicrobiaceae family, *Akkermansia* significantly increased in B6 mice after fructose treatment. In DBA feces, fructose altered the abundance of Rikenellaceae and Pseudomonadaceae. These two families were also found to be fructose responsive in B6 mice, however, the response was more dramatic in DBA mice (0.374-fold and 33.3-fold changes in DBA compared with 0.620-fold and 3.33-fold changes in B6 for Rikenellaceae and Pseudomonadaceae, respectively). No fecal microbial taxa were significantly altered by fructose in FVB mice (Table 1; **Supplemental Figure 3**A, B). In DBA cecum, family Erysipelotrichaceae and two of its genera (*Clostridum* and an unknown genus), and *Anaerostipes* were all significantly decreased by fructose. Fructose also significantly increased cecal *Bifidobacterium* in FVB mice, while no cecal taxa were significantly changed in B6 mice (Table 1; Supplemental Figure 3C, D).

**Table 1.** Differentially abundant microbiota between fructose and water groups in cecal and fecal samples of B6, DBA, and FVB mice.

| Mouse strain | Taxonomic level | Source | Fructose-responsive microbiota | Fold change |
| --- | --- | --- | --- | --- |
| DBA | Family | Cecum | Erysipelotrichaceae | 0.055* |
| B6 | Family | Feces | Rikenellaceae | 0.620* |
|  | Family | Feces | S24-7 | 1.19* |
|  | Family | Feces | Dehalobacteriaceae | 0.600* |
|  | Family | Feces | Lachnospiraceae | 0.590* |
|  | Family | Feces | Mogibacteriaceae | 0.333* |
|  | Family | Feces | Ruminococcaceae | 0.706* |
|  | Family | Feces | Turicibacteraceae | 0.348* |
|  | Family | Feces | Pseudomonadaceae | 3.33* |
|  | Family | Feces | Verrucomicrobiaceae | 7.25** |
| DBA | Family | Feces | Rikenellaceae | 0.374* |
|  | Family | Feces | Pseudomonadaceae | 33.3* |
| DBA | Genus | Cecum | *Erysipelotrichaceae (Unknown genus)* | 0.004* |
|  | Genus | Cecum | *Erysipelotrichaceae Clostridium* | 0.009** |
|  | Genus | Cecum | *Lachnospiraceae Anaerostipes* | 0.025** |
| FVB | Genus | Cecum | *Bifidobacteriaceae Bifidobacterium* | 16.5* |
| B6 | Genus | Feces | *Dehalobacteriaceae Dehalobacterium* | 0.600* |
|  | Genus | Feces | *Mogibacteriaceae (Unknown genus)* | 0.333* |
|  | Genus | Feces | *Verrucomicrobiaceae Akkermansia* | 7.27** |
| DBA | Genus | Feces | *Rikenellaceae (Unknown genus)* | 0.374* |
|  | Genus | Feces | *Pseudomonadaceae (Unknown genus)* | 60.0* |
|  | Genus | Feces | *Pseudomonadaceae Pseudomonas* | 28.0* |

<sup>1</sup>Fold change is calculated as the ratio of the relative abundance of microbiota between fructose and water groups (fructose/water). $n=8$ /group/mouse strain, \*FDR < 0.05, \*\*FDR < 0.01.

We next correlated the abundance of these fructose-responsive taxa with metabolic phenotypes in water or fructose-treated mice. In DBA mice, cecal Erysipelotrichaceae was negatively correlated with adiposity and AUC (**Supplemental Figure 4**A, B). Fecal Rikenellaceae in DBA mice had negative correlations with body weight and adiposity, and a positive correlation with AUC in the fructose group, while no significant correlation was observed with these phenotypes in the water group (**Figure 3A-F**). No phenotypic correlation was observed for the fructose-responsive taxa in B6 and FVB mice, which is not surprising given the weaker phenotypic alterations in these mouse strains in response to fructose consumption.

### Correlation of fructose-responsive microbiota with fructose signature genes in host metabolic tissues

We then analyzed the correlation between the abundance of fructose-responsive microbiota and the host fructose signature genes in liver, adipose tissue, and hypothalamus (5). We observed distinct correlation patterns in the three mouse strains: the B6 fructose-responsive taxa were correlated with only hypothalamic fructose signature genes, while the fructose-responsive taxa in DBA cecum or feces were correlated with only liver or adipose tissue signature genes, respectively (summary in **Table 2**; full list of genes correlated with fructose-responsive taxa in **Supplemental Table 1**).

**Figure 3.**
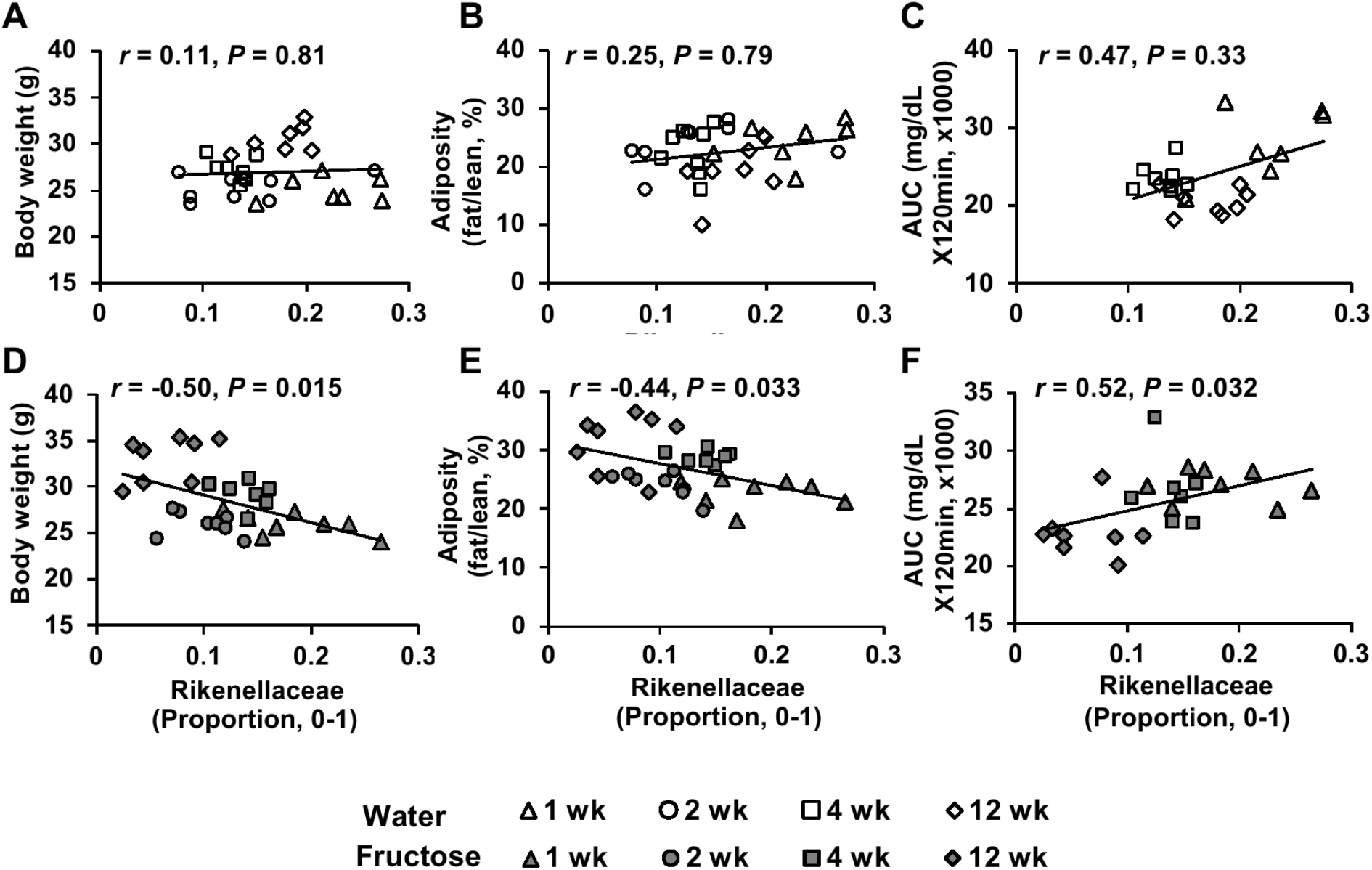
Correlation analysis of fructose-responsive microbiota with metabolic phenotypes in DBA mice. (A-C) Correlation plots between Rikenellaceae proportion and body weight (A), adiposity (B), and glucose tolerance AUC (C) across time points in the water group. (D-F) Correlation plots between Rikenellaceae proportion and body weight (D), adiposity (E), and glucose tolerance AUC (F) across time points in the fructose group. *r* = Biweight midcorrelation (*bicor*) coefficient, *P* = Benjamini-Hochberg adjusted *P*-values. *n* = 7-8/group/time point.

**Table 2.**
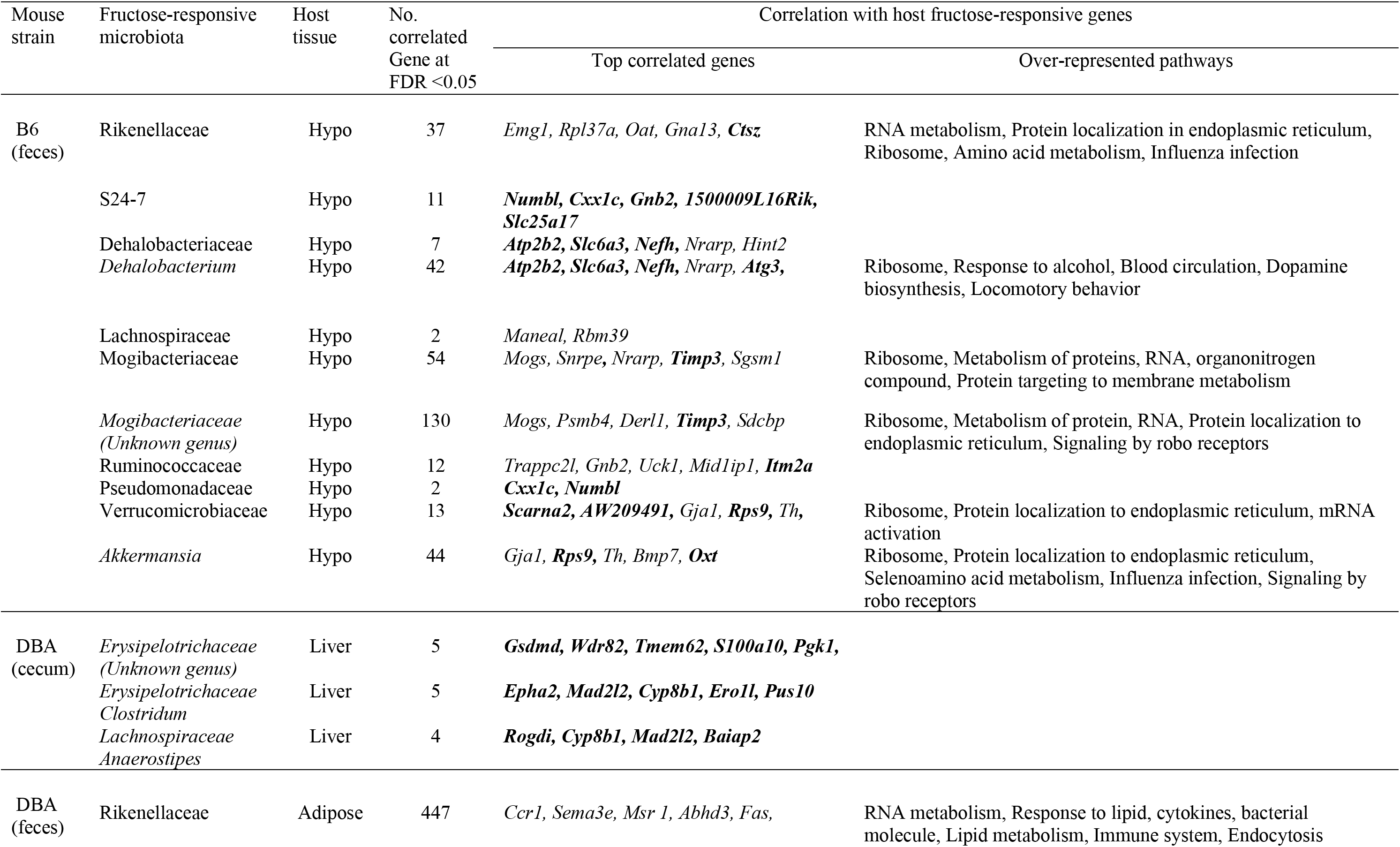
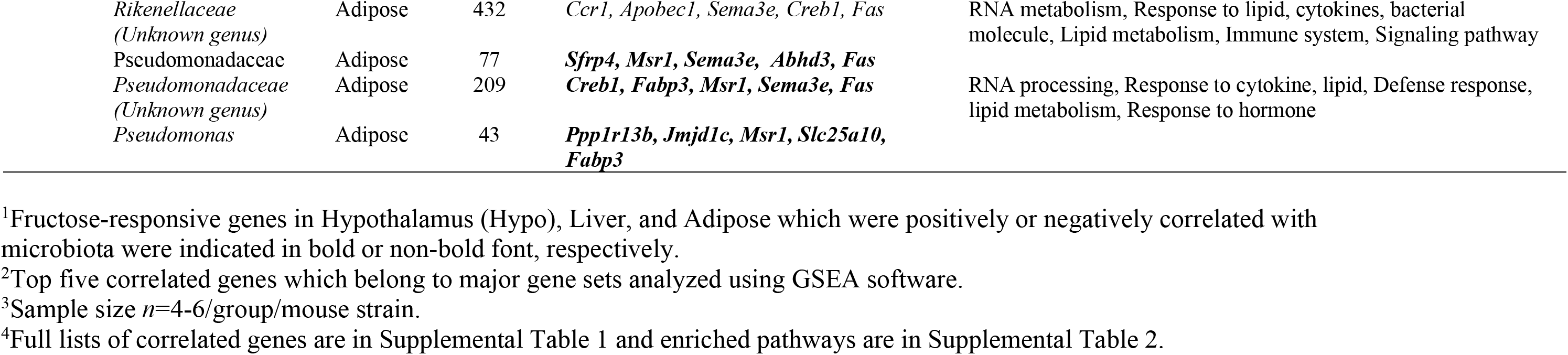
Correlation between fructose-responsive microbiota and host fructose signature genes in three metabolic tissues in B6 and DBA mice.

In B6, *Dehalobacterium* showed positive correlation with hypothalamic genes encoding the neurotransmitter transporter *Slc6a3*, a notch signaling component *Nrarp*, and an autophagy gene *Atg3*. *Akkermansia* was correlated with several neurotransmitter related genes, including *Oxt* encoding precursor of oxytocine/neurophysin 1 and *Th* encoding tyrosine hydroxylase. In DBA cecum, both *Anaerostipes* and *Clostridium* were positively correlated with *Cyp8b1* in liver, which is responsible for bile acid synthesis (34).

In DBA feces, all fructose-responsive taxa were correlated with host signature genes of the adipose tissue, and these genes were involved in lipid metabolism, immune system, response to lipid, cytokines, and hormone (**Supplemental Table 2)**. Adipose genes such as *Abhd3, Msr1, Ccr1, Creb1*, and *Fas* were correlated with Rikenellaceae and Pseudomonadaceae as well as genera within these families (Table 2; Supplemental Table 2). Taken together, these correlations suggest that gut microbiota may interact with host genes in a mouse strain- and tissue-specific manner in response to fructose.

### Alteration of gut microbiota modulates fructose response

Since B6 and DBA mice showed disparate metabolic responses to fructose, we tested whether B6 microbiota confers resistance and DBA microbiota confers vulnerability to fructose effects by transplanting B6 feces to antibiotic-treated DBA mice and vice versa (**Figure 4A**). Using 16S rDNA sequencing, we confirmed that the recipient mice gut microbiome shifted after fecal transplant (**Supplemental Figure 5**A, B).

**Figure 4.**
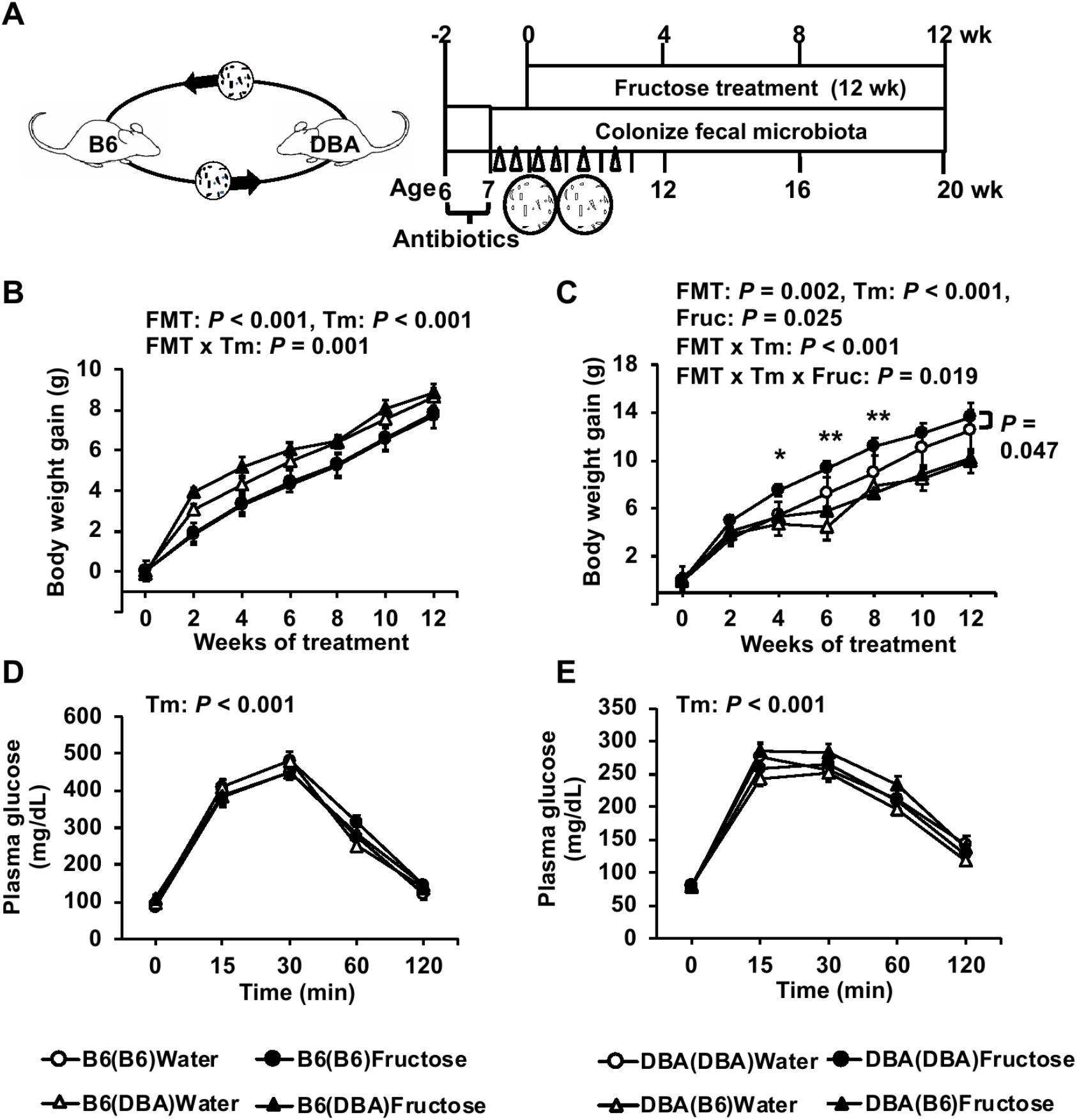
Metabolic phenotypes post fecal transplant in B6 and DBA mice with or without fructose consumption. (A) Schematic design of fecal microbiota transplant (FMT). (B-E) Body weight gain (B, C) and glucose tolerance (D, E) of recipient B6 and DBA mice, respectively, with or without 8% fructose water. Data are presented as means ± SEM, *n* = 7-14/group. The *P*-values of the main factors (FMT, fructose, time) and interactions by three-way repeated-measures ANOVA are shown on the top of the graph. Asterisks in (C) indicate time points at which significant differences were found between DBA(DBA) and DBA(B6) under fructose treatment based on two-way repeated-measures ANOVA with Sidak’s post hoc test; \**P* < 0.05, \*\**P* < 0.01. Fructose effects within each FMT group across time points were assessed by two-way repeated-measures ANOVA, and significant difference between fructose and water treatments for the DBA(DBA) FMT group is indicated by a *P*-value with a side bar (C). B6(B6): B6 mice receiving B6 feces; B6(DBA): B6 mice receiving DBA feces; DBA(B6): DBA mice receiving B6 feces; DBA(DBA): DBA mice receiving DBA feces; Fruc: Fructose; Tm: Time.

When the main effects of FMT, fructose, and time were tested, there were significant FMT effects on weight gain in both B6 (*P* < 0.001, Figure 4B) and DBA mice (*P* = 0.002 Figure 4C), but there was no effect of FMT on glucose tolerance in both mouse strains (Figure 4D, E). Overall, there was a significant fructose effect on weight gain in DBA mice (*P* = 0.025; Figure 4C), which was not observed in B6 mice (Figure 4B). In those fed fructose, body weight gain in DBA mice receiving B6 bacteria [DBA(B6)] was significantly lower compared to DBA(DBA) at 4 weeks (*P* = 0.028), 6 weeks (*P* = 0.006), and 8 weeks (*P* = 0.007). In contrast, there was no significant fructose effect in B6 mice receiving DBA bacteria [B6(DBA)] or in B6(B6) mice (Figure 4B). These results suggest that DBA microbiota failed to induce fructose sensitivity in B6 mice. On the other hand, DBA(B6) mice no longer displayed fructose-induced weight gain (fructose effect *P* = 0.66) as seen in DBA(DBA) mice (fructose effect *P* = 0.047, Figure 4C), supporting that B6 microbiota conferred resistance to body weight gain upon fructose consumption.

The result from FMT experiment supports a causal role of B6 microbiota in conferring fructose resistance to DBA. We next focused on prioritizing the potential microbes in B6 that may determine the fructose resistance phenotype. *Akkermansia* was found to be a highly plausible candidate to explain the dampened response to fructose in B6 for the following reasons. First, *Akkermansia* has been previously demonstrated to carry anti-obesity and insulin sensitizing effects (32,35,36). The beneficial effect of *Akkermansia* was also previously observed in mice fed a high-fat/high-sucrose diet (37). Secondly, it was depleted in the vulnerable strain DBA but was highly abundant in both resistant mouse strains, B6 and FVB (Figure 2G). Thirdly, fructose treatment caused an increase in *Akkermansia* in B6 mice (Table 1; Supplemental Figure 3A). Lastly, *Akkermansia* abundance in B6 is significantly correlated with a large number of hypothalamic genes such as *Oxt* and *Th* that regulate metabolism (Table 2; Supplemental Table 1).

To test the role of *Akkermansia* in protecting against fructose-induced metabolic dysregulation as predicted above, we gavaged DBA mice with *Akkermansia muciniphila* (AM) or PBS media along with fructose treatment for 8 weeks (**Figure 5A**). There were significant effects of AM on both body weight gain (*P* < 0.001) and glucose tolerance (*P* < 0.001) (Figure 5B, C). Under fructose treatment, DBA mice receiving AM had significantly lower weight gains at 3 (*P* = 0.023) and 5-8 weeks (*P* < 0.01) compared to control mice receiving PBS. No AM effect was observed in the water treatment mice (*P* = 0.14). Within AM or PBS group, fructose had significant effects on body weight gain in the PBS control group (*P* = 0.037), whereas DBA mice receiving AM no longer displayed fructose-induced body weight gain as seen in control mice (Figure 5B). Furthermore, fructose increased glucose intolerance in PBS control group (fructose effect *P* = 0.044), whereas AM treatment abrogated fructose-induced glucose intolerance in DBA mice (Figure 5C). These results support that *Akkermansia* confers resistance to fructose-mediated metabolic dysregulation.

**Figure 5.**
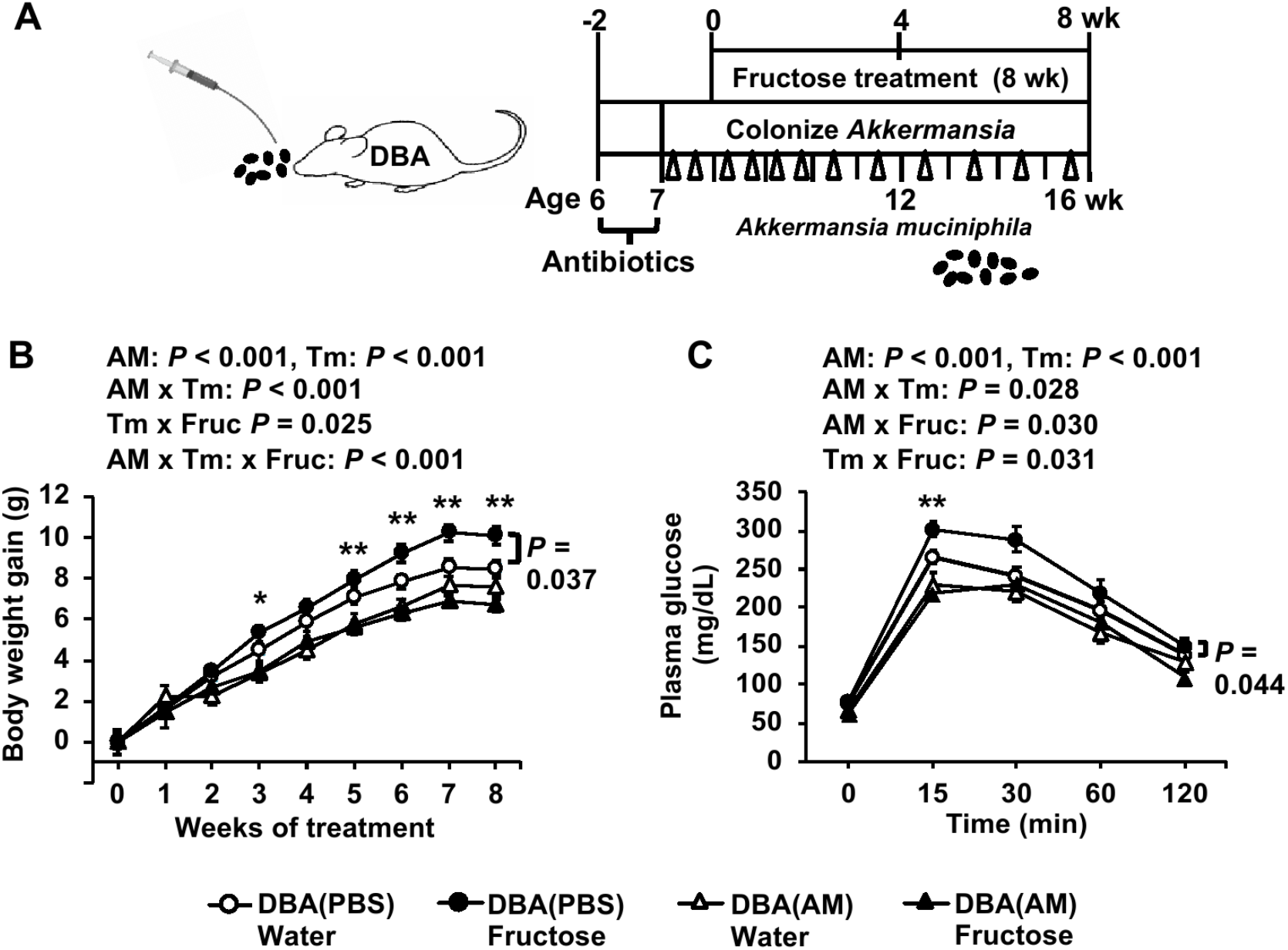
Metabolic phenotypes post *Akkermansia* colonization in DBA mice with or without fructose consumption. (A) Schematic design of *Akkermansia muciniphila* (AM) colonization. PBS serves as the control for AM. (B-C) Body weight gain (B) and glucose tolerance (C) of recipient DBA mice with or without 8% fructose water. Data are presented as means ± SEM, *n* = 8-14/group. The *P*-values of main factors (AM, fructose, time) and interactions by three-way repeated-measures ANOVA are shown on the top of the graph. Asterisks indicate time points at which significant differences were found between the AM colonization and PBS control groups under fructose treatment based on two-way repeated-measures ANOVA with Sidak’s post hoc test; \**P* < 0.05, \*\**P* < 0.01. Fructose effects within PBS or AM group across time points were assessed by two-way repeated-measures ANOVA. A significant fructose effect on weight gain and glucose tolerance in mice receiving PBS is indicated by a *P*-values with a side bar (B and C). Fruc: Fructose; Tm: Time.

## Discussion

Our previous study showed that three mouse strains representing a range of genetic diversity differed in their metabolic and transcriptomic responses to high fructose treatment (5). Since gut microbiota is an important modulator of metabolic capacity (7), here we tested the hypothesis that disparate fructose responses among mouse strains were at least partially driven by the gut microbiota. Our 16S rDNA sequencing analysis revealed that baseline microbiota composition and its response to fructose varied by mouse strains. The fecal transplant data indicates that B6 mice carry microbiota that confer resistance to fructose-induced body weight gain. We next evaluated candidate taxa to explain the dampened response to fructose in B6 mice. We prioritized *Akkermansia* since it is enriched in B6 compared to DBA mice, which have lower levels. Indeed, gavaging *Akkermansia muciniphila* to DBA mice mitigated fructose-induced obesity and glucose intolerance. These results support a causal role of gut microbiota in determining the differential metabolic responses to fructose among genetically diverse mouse strains.

As initial colonizing microbial species are important for establishing a favorable environment for bacterial growth in a particular context (38), differences in baseline microbial composition between mouse strains can lead to variations in the metabolic processes of individuals in response to diet (31,32). We found that *Akkermansia*, *Turicibacter*, and S24-7 were lower in DBA mice but were abundant in B6 and FVB mice. On the other hand, the baseline levels of *Lactobacillus*, Clostridiales and Lachnospiraceae were higher in DBA mice. Previously, *Akkermansia* has been associated with obesity resistance and improved metabolic parameters in humans and mice, and beneficial effects of dietary interventions have been associated with the higher abundance of *Akkermansia* at baseline (32,36). S24–7 has protective association against diabetes, whereas Lachnospiraceae promotes pathogenesis of diabetes in NOD mice (39). The observed lack of protective *Akkermansia* and S-24, along with the higher abundance in pathogenic Lachnospiraceae in DBA mice, agrees with the vulnerability of DBA mice to fructose-induced metabolic dysregulation. Therefore, these bacterial taxa that differ significantly at baseline between strains have the potential to regulate the differential response to fructose treatment.

In addition to differences in baseline microbes that can explain inter-mouse strain variability, bacteria altered by fructose may also play a role in the variability in metabolic responses to fructose. It has been suggested that fructose shifts the gut microbiota and leads to a westernized microbiome acquisition with altered metabolic capacity, resulting in development of obesity or metabolic disorders (40). In our study, we found various microbes with significantly altered abundances in DBA and B6 but not FVB, which may relate to their differential sensitivity to fructose. Cecal Erypsipelotricaceae and *Anaerostipes* in DBA mice decreased in abundance upon fructose consumption (Table 1; Supplemental Figure 3), and they are known as butyrate-producing bacteria (41). Butyrate promotes the intestinal barrier development, and decreased butyrate production can increase intestinal permeability (42). In both the B6 and DBA feces, we also detected decreased abundance of Rikenellaceae and increased abundance of Pseudomonadaceae following fructose treatment (Table 1). Increases in Pseudomonadaceae and *Pseudomonas* were previously found in obese individuals with higher insulin resistance (43). Therefore, despite the numerous differences between B6 and DBA, there were shared microbial changes in response to fructose consumption. The weaker phenotypic responses in B6 can be a result of compensatory balancing effects of higher abundances of beneficial bacteria such as *Akkermansia* and S24-7 in B6.

We detected more fructose-responsive microbiota in B6 mice than in DBA and FVB mice (Table 1), and, interestingly, most of these B6 taxa were associated with the hypothalamic fructose signature genes (Table 2; Supplemental Table 1). Among them are genes that are important for energy homeostasis such as *Nrarp, Qxt*, and *Th*. *Nrarp* encodes an intracellular component of the Notch signaling pathway and regulates differentiation of mouse hypothalamic arcuate neurons responsible for feeding and energy balance. Dysregulation of this homeostatic mediator is an underlying cause of various diseases ranging from growth failure to obesity (44). *Qxt* encodes oxytocin which is the anorexigenic peptide. Oxytocin maintains homeostasis in feeding-related behavior (45). *Th* gene encodes tyrosine hydrogenase (TH). Hypothalamic arcuate nucleus TH neurons play a role in energy homeostasis, and silencing of TH neurons reduces body weight (46). Our previous mice study (2) reported that fructose is an inducer of both genomic and epigenomic variability in hypothalamus and has the ability to reorganize gene networks that plays a central role in metabolic regulation and neuronal processes. The current study suggests that fructose-induced transcriptomic changes in the hypothalamus of B6 mice could be partly driven by fructose-responsive gut microbiota.

In contrast to B6 mice, fructose-responsive microbiota in DBA mice were associated with lipid and inflammatory genes in the adipose tissue (Table 2; Supplemental Table 2). *Abhd3* is a crucial factor for insulin resistance in adipose tissue (47); *Sema3e* contributes to inflammation and insulin resistance in obese mice (48); *Msr1* is macrophage scavenger receptor 1 and provides protection from excessive insulin resistance in obese mice (49); *Creb1* promotes expression of transcriptional factors of adipogenesis and insulin resistance in obesity (50); *Fas* contributes to adipose tissue inflammation, hepatic steatosis, and insulin resistance induced by obesity (51). Therefore, these adipose-tissue genes correlating with fructose-responsive bacteria in DBA are relevant to the increased adiposity and compromised insulin sensitivity seen in DBA mice. However, we acknowledge that the correlative relationship observed here does not directly imply causation, and future experiments are needed to directly test the causal role of the genes as well as the bacteria implicated.

Our fecal transplant study support that B6 mice carry gut microbes that confer resistance to fructose-induced metabolic syndrome, while DBA microbiota did not significantly induce fructose sensitivity in B6 mice. We further demonstrate that *Akkermansia* partially mediates the protective effect of B6 microbiota. This is the first time that *Akkermansia* is implicated in determining fructose response. Given the recognized therapeutic potential of modulating the gut microbiota (8), probiotic treatment with *Akkermansia* may represent a viable approach to mitigate fructose-induced metabolic abnormalities. In addition to *Akkermansia*, other bacteria may also play roles in modulating the differential fructose response between individuals. Future efforts will explore the pathogenic or protective roles of other bacterial species, such as *Lactobacillus*, *Turicibacter*, or *Pseudomonas*.

In summary, our multi-strain, multi-omics (gut microbiome, transcriptome, and phenome) integrative studies of inter-individual variability in fructose-induced metabolic syndrome established a causal role of gut microbiota in modulating host responses to fructose, a significant metabolic risk in modern societies. Importantly, our study identified key microbial species that may serve as preventative or therapeutic targets for metabolic syndrome. By exploring gut-host interactions, our study also opens numerous hypotheses regarding specific microbiota-host interactions to be further elucidated in the future.

## Acknowledgements

The authors thank Dr. Richard C. Davis for assistance of oral gavage and fecal transplant, Dr. Yuqi Zhao and Dr. Zeyneb Kurt for the assistance with data analysis, and Dr. Sana Majid, Paul Patel, Maya Singh, Jessica Yang and Justin Yoon for helping with animal experiments.

XY, EYH, ISA, and CO designed the research; ISA, CAO, JML, ZY, IC, GD, PC, HRB and JD conducted research including animal experiments; JML, ISA, GZ, IK, JD analyzed data; ISA, XY and JML wrote the manuscript; FG-P, AL and all authors contributed to manuscript revision; XY had primary responsibility for the final content; All authors read and approved the final version of the paper.

