## Supplemental Figures for "Host Genetic Background and Gut Microbiota Contribute to Differential Metabolic Responses to Fructose Consumption in Mice"

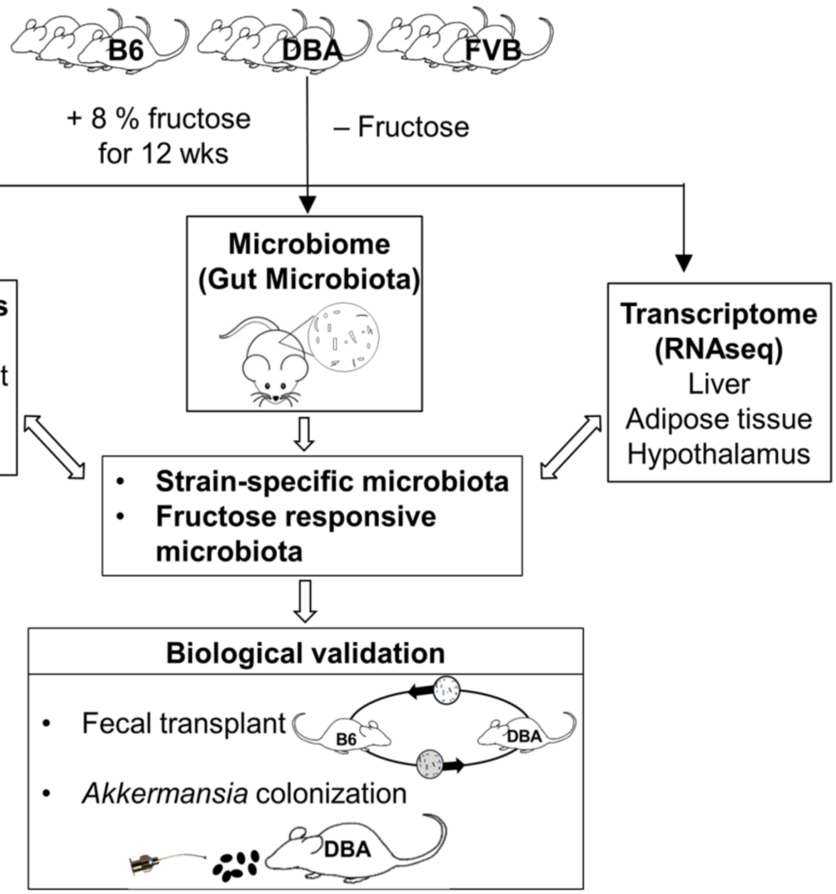


**Supplemental Figure 1. Overall study design.** Three mouse strains, C57BL/6J (B6), DBA/2J (DBA), FVB/NJ (FVB), were used to study the differential metabolic responses to fructose treatment. We treated mice with 8 % fructose water for 12 weeks and collected metabolic phenotypes, microbiome data, and transcriptome data from host individual tissues (hypothalamus, liver, and mesenteric adipose tissue). The gut microbiota showing differential baseline levels between mouse strains as well as fructose-responsive microbiota were correlated with metabolic phenotypes and fructose signature genes in individual tissues to prioritize microbial species associated with host responses. The causal role of gut microbiota was validated by fecal transplant between B6 mice (fructose resistant) and DBA mice (fructose sensitive). Finally, *Akkermansia* was inoculated to DBA mice to determine its causal role in mitigating the fructose response in DBA mice.

### Supplementary Data

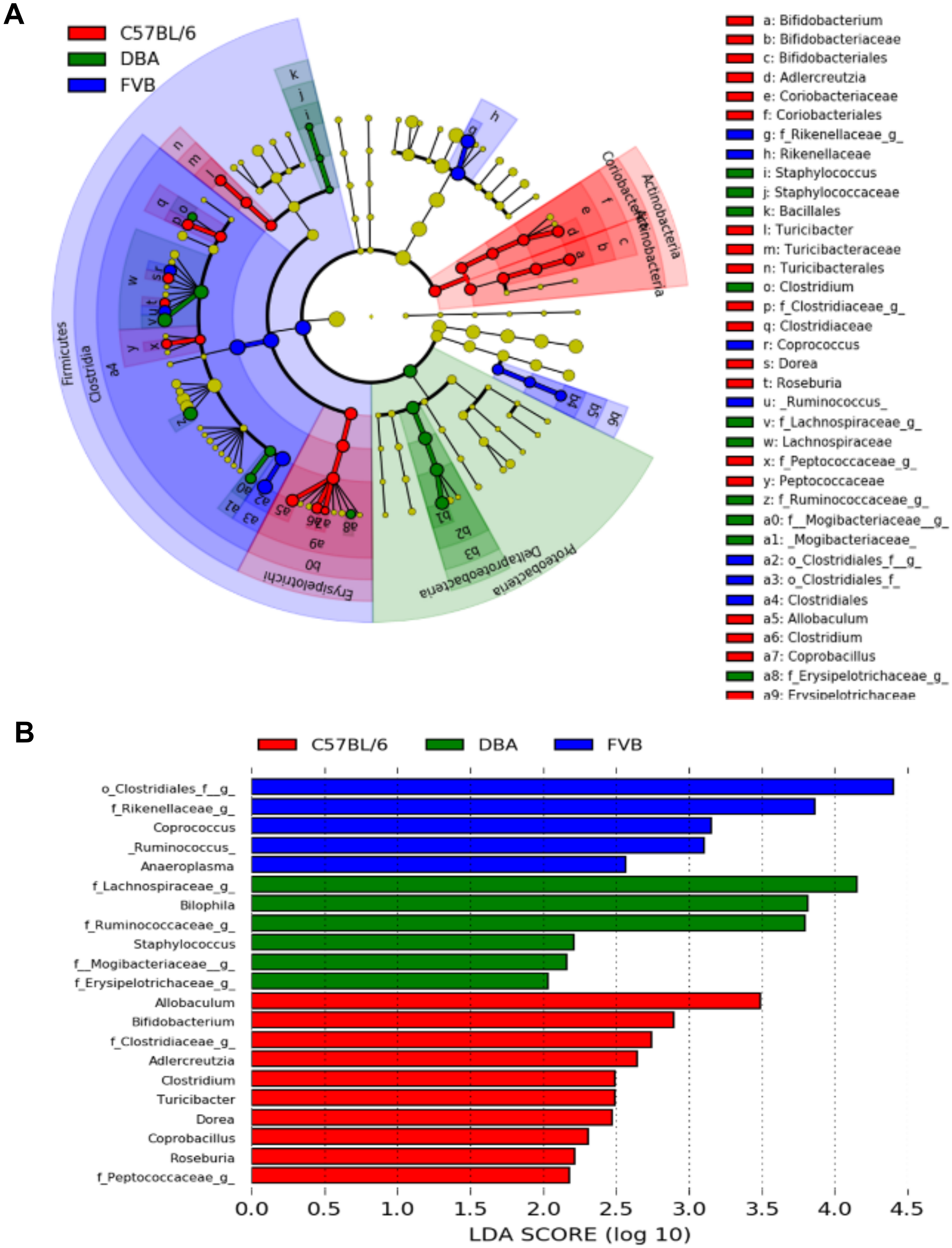

### Supplementary Data

**C**

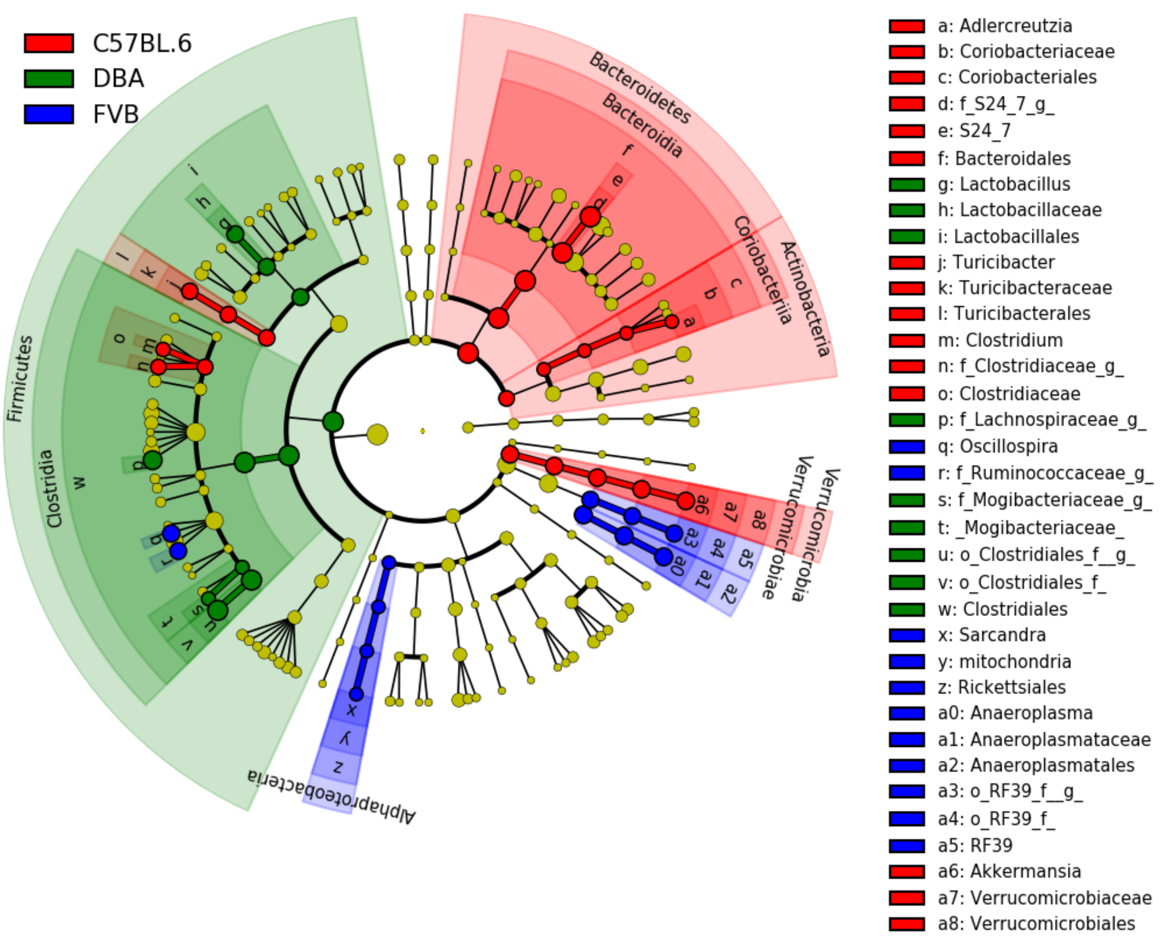


**D**

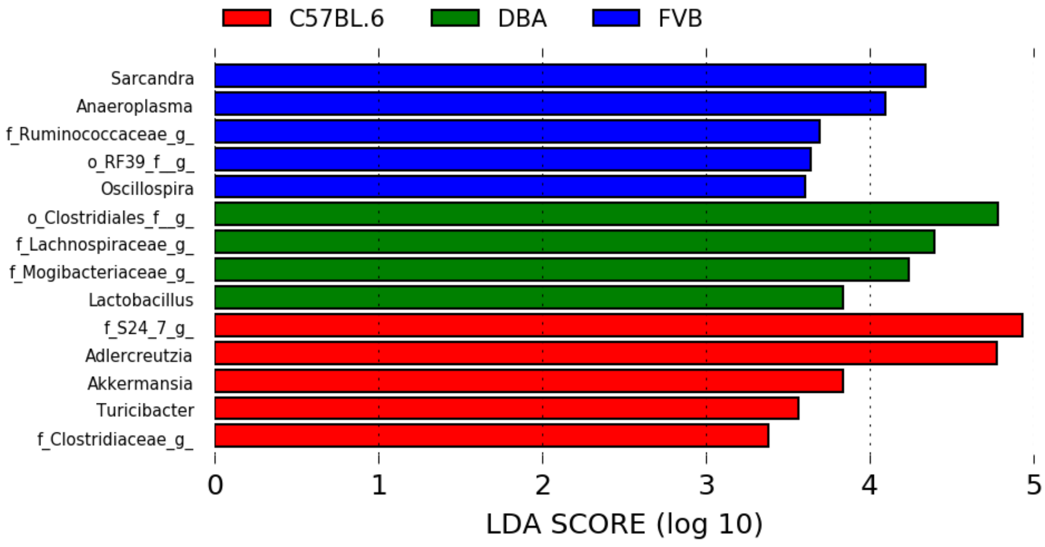

#### Supplementary Data

**Supplemental Figure 2. Identification of differentially abundant baseline microbiota between B6, DBA, and FVB mice at baseline.** Linear discriminant analysis (LDA) effect size (LEfSe) was used to identify taxa that discriminated between the three mouse strains using standard parameters ($p < 0.05$, LDA score > 2.0). The baseline taxa levels were analyzed using the microbiota from mice of the water group at all time points. Cladogram shows the significant taxa at all taxa levels (colored circles) of cecal (A) and fecal (C) microbiota of three strains, but they are only labeled to the family level. Significantly different genera in each mouse strain of cecal (B) or fecal (D) microbiota are shown in the bar graphs. Each ring of the cladogram represents a different taxonomic level, starting with kingdom in the center and ending with genus in the outer ring. Colors indicate the mouse group (red, green, blue for B6, DBA, and FVB, respectively) with the highest mean of differential features for which significant differences between strains were found. Taxa (yellow) that do not show significant difference in abundance between groups are unlabeled. $n$ = 8/group.

#### Supplementary Data

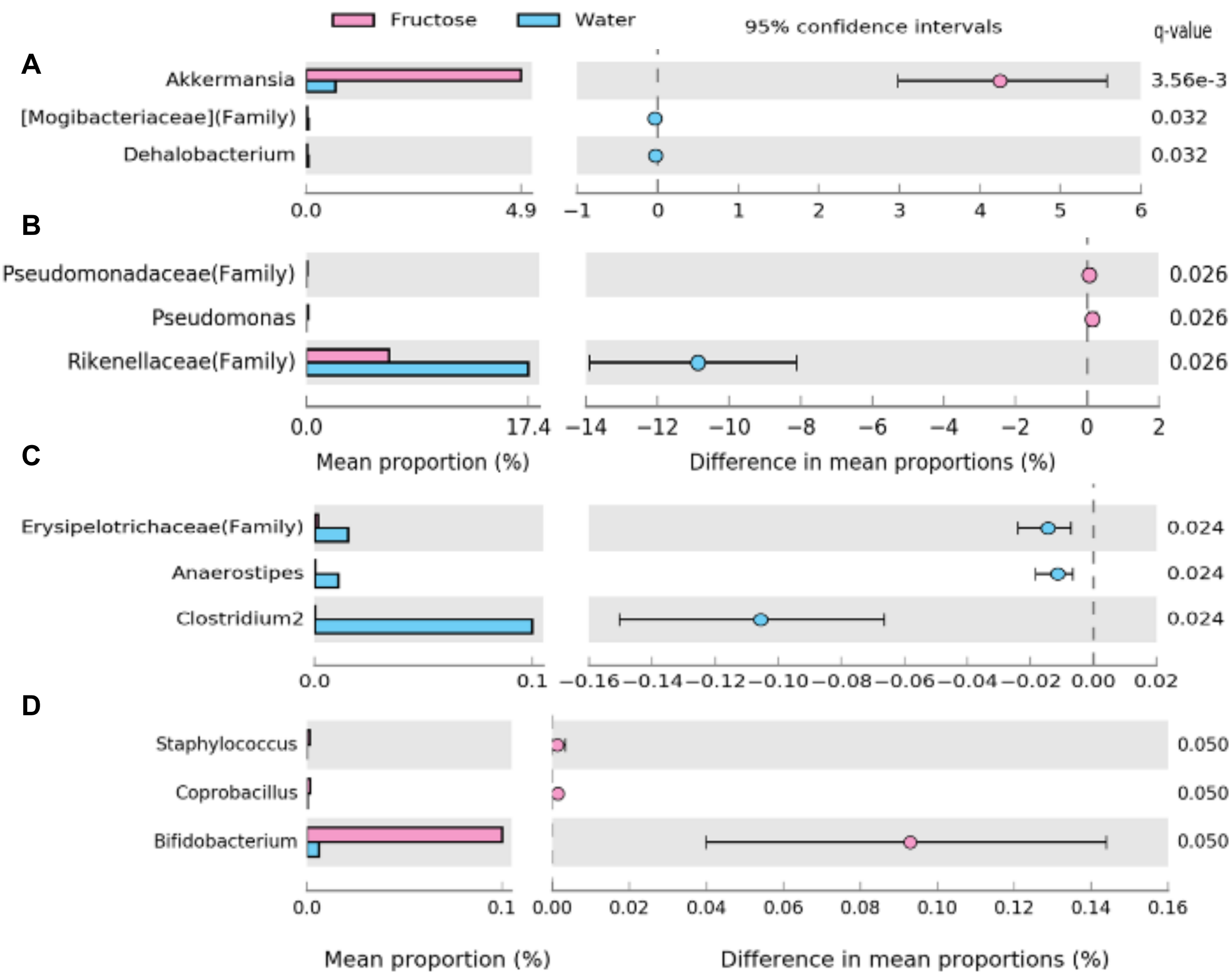


**Supplemental Figure 3. Differentially abundant gut microbiota between fructose and water groups in B6, DBA, and FVB mice.** Mean proportions of microbiota in water (blue) and 8 % fructose (pink) groups are plotted. The difference in mean proportions were visualized using extended error bar plot. (A) B6 fecal, (B) DBA fecal, (C) DBA cecal, (D) FVB cecal bacteria. Multiple testing corrected q values are shown to the right of each plot. $n$ = 8/group.

**Supplementary Data**

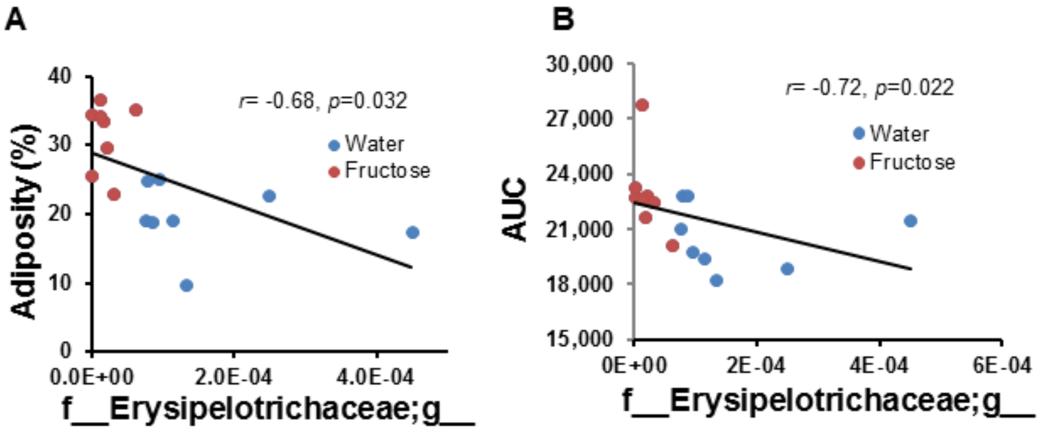


**Supplemental Figure 4. Correlation between fructose-responsive cecal microbiota in DBA mice and metabolic phenotypes.** The proportion of the fructose-responsive cecal microbiota at genus level was correlated with adiposity (A) and AUC (B). Data were collected for both water and fructose group at 12 weeks of fructose treatment. *r* = Biweight midcorrelation (*bicor*) coefficient, *P* = Benjamini-Hochberg adjusted *P*-values. *n* = 7-8/group.

**Supplementary Data**

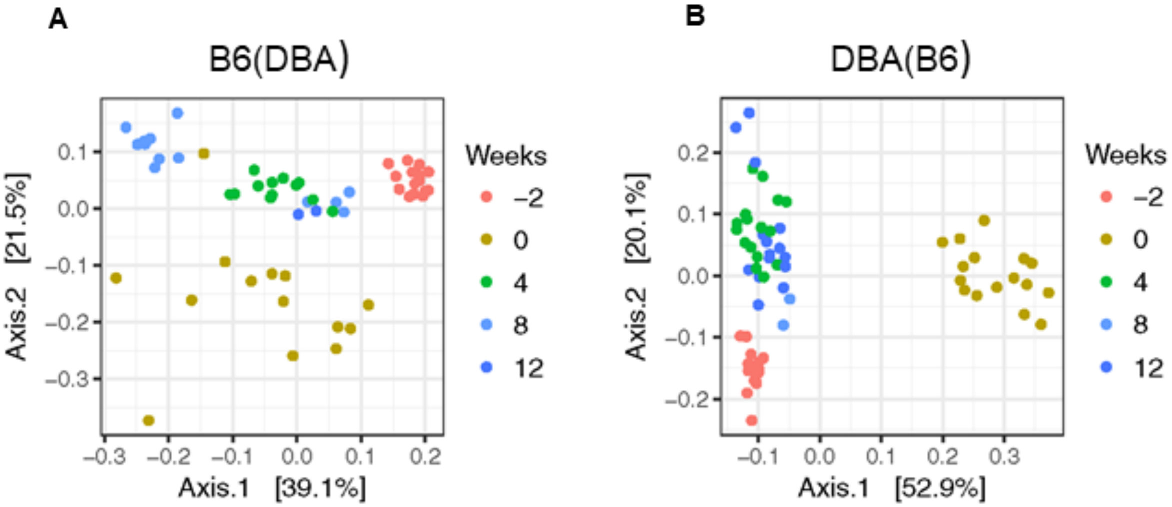


**Supplemental Figure 5. Principal coordinate analysis (PCoA) of fecal microbiota in the fecal transplanted B6 and DBA mice.** 6-week-old recipient mice were treated with antibiotics for one week, and then fecal supernatant from donor B6 mice was orally transplanted to the recipient DBA mice, and vice versa. B6 mice that received DBA feces are designated B6(DBA) (A), and DBA mice that received B6 feces are indicated as DBA(B6) (B). After 1 week of fecal transplant, B6(DBA) and DBA(B6) mice were challenged by 8 % fructose water or water control for 12 weeks. Fecal samples were collected at weeks -2, 0, 4, 8, 12 of fructose treatment. The start of the antibiotic treatment occurred at -1 weeks. *n* = 13-16.
